# A mechanically stable neural probe for percutaneous high-resolution, multi-channel recordings in peripheral nerves

**DOI:** 10.64898/2026.04.10.717712

**Authors:** Sofiia Demchenko, Erkan Yilmaz, Anoushka Jain, Stas Koulchitsky, James P. Dunham, Anthony E. Pickering, Barbara Namer, Simon Musall, Viviana Rincón Montes

## Abstract

The development of neural probes has enabled a deeper understanding and improved treatment for neurological disorders. Microneurography is currently the gold standard for assessing the electrophysiological signature of pain mechanisms in the human peripheral nervous system. However, its clinical utility is limited by the low recording yield and signal-to-noise ratio of single-electrode probes. To overcome these limitations, we developed mechanically robust, multi-electrode probes designed for acute percutaneous insertion and recording in peripheral nerves. The electrical and mechanical stability of these probes was confirmed through repeated insertions in artificial human skin and rat peripheral nerves. In addition, *ex vivo* and *in vivo* experiments demonstrated enhanced functional performance, with multi-site recordings enabling the isolation of single-fiber activity. Importantly, our probes can be operated analogously to conventional microneurography needles while substantially increasing the information yield, providing enhanced capabilities for minimally invasive peripheral nerve assessment.

## Introduction

Chronic pain is a major public health issue that substantially impairs quality of life, not only due to persistent aversive sensory symptoms but also through its profound psychological and social consequences. A major subset of chronic pain is neuropathic pain due to lesions or diseases of the somatosensory nervous system, affecting approximately 10% of the global population ^1^. The dysfunction or degeneration of peripheral nerve fibers is a particularly prominent cause of neuropathic pain, leading to ongoing pain, hyperalgesia, and allodynia in affected patients while being particularly difficult to diagnose and treat ^1,2^. For example, small-fiber neuropathy (SFN) involves the thinly myelinated Aδ fibers and unmyelinated C fibers, which are spatially intermixed within peripheral nerves and mediate pain (*e.g.*, heat, pressure, or chemical stimuli), thermal sensitivity, affective touch, and autonomic functions ^3–6^. Among these, nociceptors are a specialized subset of sensory fibers that detect and encode noxious stimuli, initiating the neural activity that underlies pain perception. Importantly, distinct subtypes of peripheral nerve fibers also display variable receptive fields, response dynamics, and conduction velocities, which makes it challenging to resolve their individual activity using compound nerve recordings. To objectively characterize the physiological phenotype of SFN, and evaluate the efficacy of emerging treatments, it is therefore necessary to record and analyze the spiking activity of individual nerve fibers ^7–9^. This requires single-unit resolution techniques that can capture the diverse pathological signatures of small fiber dysfunction, which are often missed by standard clinical tests such as skin biopsies or nerve conduction studies ^10^. Peripheral nerve recordings provide a powerful means to directly measure action potentials in afferent fibers within their native tissue environment, offering a unique opportunity to assess the conduction properties and firing pattern of individual fibers ^11^.

For human subjects, microneurography remains the only technique currently available to record action potentials from individual axonal fibers in the peripheral nervous system. This method uses a single-electrode, tungsten needle to record the action potentials of axons and has been instrumental in functionally characterizing different subtypes of nerve fibers ^7,12^. For example, activity-dependent conduction velocity slowing in nociceptors can be used to distinguish different functional subclasses and link them to specific pathological states ^13^. Microneurography can also capture the activity underlying complex sensations like itch-related discharge pattern encoded signaling, detect age-related changes in nerve function, and differentiate types of neuropathies based on patterns of abnormal nociceptor activity, such as ongoing spontaneous firing or evidence of denervation ^14,15^. These insights are critical for translational research linking animal models of neuropathy to human pathophysiology and offer the means to identify physiological biomarkers for diagnosing SFN and monitoring therapeutic outcomes in human patients ^9,16–18^.

However, despite its advantages, microneurography is limited by its reliance on single-electrode recordings. This constrains the number of fibers that can be recorded per insertion and increases the burden on patients, as identifying individual nerve fibers is often time-consuming. Moreover, simultaneously recorded fibers with comparable tuning can be difficult to disambiguate since the electrical signals are superimposed on a recording channel. In contrast, implantable micro-electrode arrays (MEAs), such as high-density silicon probes containing multiple electrodes within the same shank, have been widely adopted in central nervous system recordings due to their ability to simultaneously detect activity from multiple neural units ^19–21^. MEAs also enable improved clustering and classification of action potentials to single units originating from different sources based on spatial and temporal features, significantly increasing data yield while reducing experiment duration ^20,22,23^. Recently, silicon-based MEAs were also used in peripheral nerve recordings in rat models, demonstrating their potential for recording spontaneous and evoked activity from a large number of axonal nerve fibers *in vivo* ^4^. Yet, technical challenges remain. Existing silicon-based MEA probes, although capable of penetrating peripheral nerves, are very fragile and unsuited for insertions through mechanically resistant tissues, such as the skin and connective tissue. The brittleness of silicon-based probes also poses significant risks in clinical settings, as breakage within a nerve could result in lasting damage. Conversely, flexible MEAs used in the central nervous system could mitigate the risk of mechanical breakage but are not mechanically stable enough for the insertion and retrieval requirements of an acute percutaneous implantation. Existing MEAs are therefore unsuitable for acute multichannel recordings of peripheral nerves in human subjects.

To bridge this gap, we present an enhanced microneurography probe that integrates flexible MEAs with microneurography-inspired needles, designed for acute peripheral nerve recordings with improved mechanical stability. The resulting peripheral nerve probe (PNP) allows for percutaneous insertion of a 32-channel recording array while maintaining a minimal cross-sectional footprint comparable to a traditional microneurography needle. Here, we describe the development of a PNP and present its mechanical and electrochemical characterization, together with robustness and functional validation *in vitro* and *in vivo* using artificial human skin (AHS) and rodent models. The dense electrode arrangement of the PNP enables simultaneous and independent recording of axonal signals from multiple sites within the peripheral nerve, therefore increasing the possible yield from acute nerve recordings while reducing the localization time. Multi-site recordings also facilitate spike sorting based on the spatial distribution of action potentials, aiding in the separation of functionally distinct fibers. Critically, the PNP contains no brittle materials such as silicon or glass, reducing the risk of nerve damage and making it generally suitable for acute recordings in human subjects. Overall, this design provides a new platform for acute recordings in neural tissues with substantial mechanical support, such as peripheral nerves or the spinal cord, and establishes a technological basis for future improvements in SFN diagnostics and neuropathic pain research.

## Results and discussion

### Design and fabrication of the PNP

PNPs need to combine both flexibility and sufficient mechanical stability for percutaneous insertion and acute recordings in peripheral nerves, such as the superficial peroneal in humans. Our PNP design is based on a thin and flexible parylene-C (PaC) based micro-electrode array (flexMEA) that is bonded to a carrier needle with UV-curable medical adhesive (Fig. 1A). Due to their low cost and intended use for skin penetration, commercially available steel acupuncture needles were selected as carrier needles. To minimize insertion-related trauma, needles with a shank diameter of 120 µm were chosen, ensuring that the nerve-penetrating section was substantially smaller than the targeted nerve. In humans, the diameter of the superficial peroneal nerve is typically in the range of a few millimeters ^24^, whereas in small laboratory animals such as rats, superficial nerves such as the saphenous nerve is approximately 300 µm ^25^. In comparison, typical microneurography needle probes, which have been shown to be mechanically robust, usually exhibit shank diameters around 200 µm ^26^.

**Figure 1.**
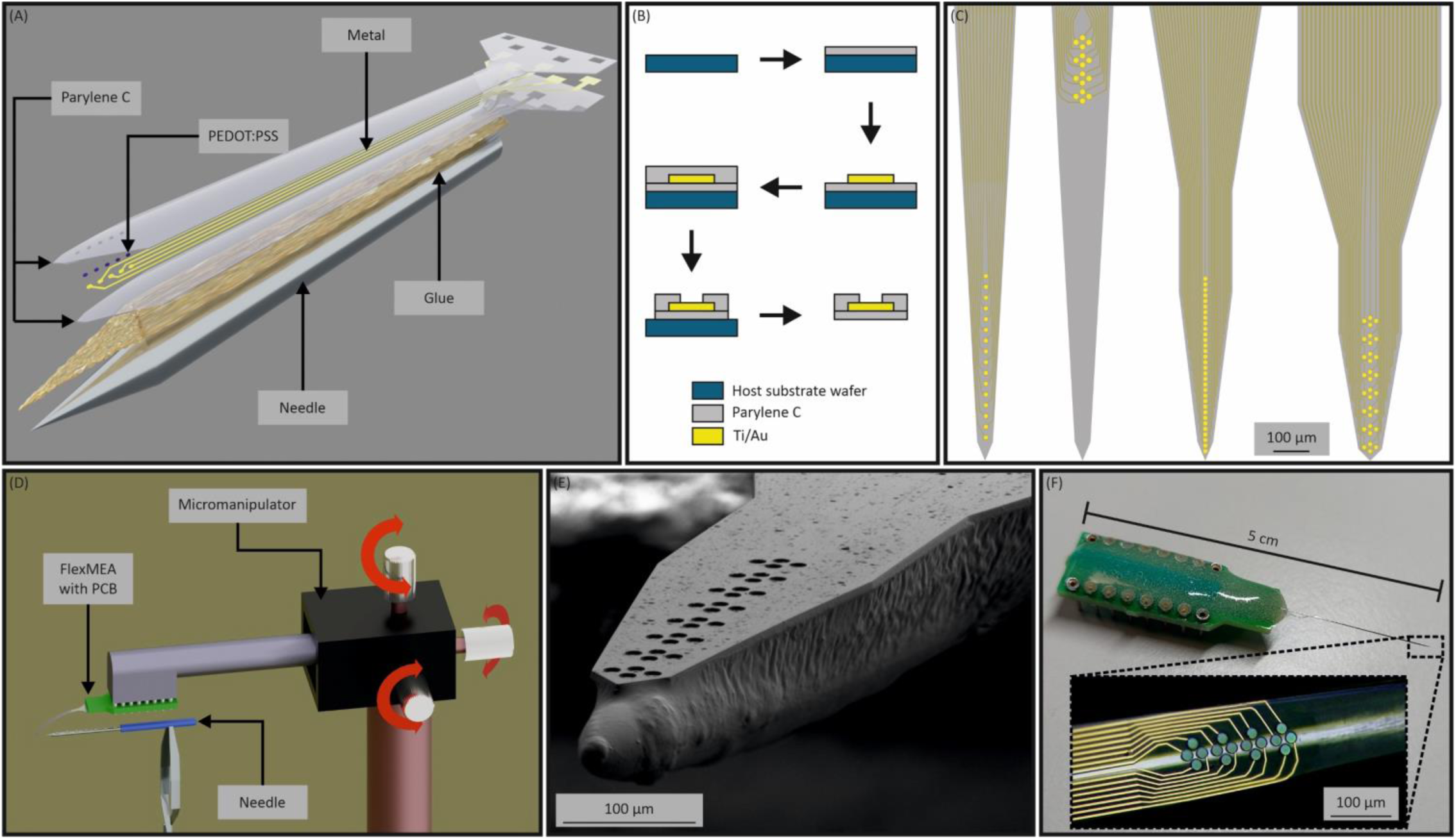
Design and fabrication of the PNP. **(A)** Exploded-view drawing of a PNP without PCB and connector. Shown are the needle with glue layer and above the first PaC layer, metal layer, and second PaC layer of the flexMEA. **(B)** Schematic of the PNP microfabrication steps. PaC deposition on a substrate wafer was followed by a structured titanium and gold metal layer for electrodes and feed lines, deposition of second PaC layer, and etch of shape and passivation opening. **(C)** Overview of different electrode arrangements, from left to right: 16 electrodes in line, 16 electrodes in tetrode, 32 electrodes in line, and 32 electrodes in tetrode arrangement, respectively. **(D)** Schematic of the needle-MEA-bonding setup to position the flexMEA on the needle carrier with a micromanipulator. **(E)** Scanning electron microscope (SEM) picture of an exemplary 32-channel PNP. **F)** Example of a finalized PNP with PCB to connect to an electrophysiological recording system; inset: exemplary image of a 16-channel PNP with tetrode arrangement and electrodeposited PEDOT:PSS.

During the exploratory phase of the study, a substantial number of detachments of the flexMEA from the metallic needles were observed. To address this issue, the metallic carrier needles were first coated with a 3 µm layer of PaC to improve the overall adhesion between the flexMEA and the carrier needle. The flexMEAs were fabricated using standard cleanroom microfabrication techniques ^27,28^. First, a 3-6 µm thick PaC layer was deposited on a host substrate wafer, followed by the implementation of metallic electrodes, feedlines, and contact pads. These conductive paths were subsequently insulated with a second PaC layer of the same initial thickness, after which the electrodes and contact pads, as well as the overall shape of the flexMEAs were defined by reactive ion etching. Finally, flexMEAs were released from the host substrate wafer (Fig. 1B).

We designed flexMEAs with different geometries and electrode counts, including both linear and tetrode electrode arrangements with a total of 16 or 32 electrodes (Fig. 1C). The linear arrangement comprised 12 µm-diameter electrodes spaced evenly with a pitch of 30 µm and 17 µm along the longitudinal axis of the flexMEA for 16- and 32-channel PNPs, respectively. In the tetrode configuration, the 15 µm- and 12 µm-diameter electrodes were grouped in diamond-like clusters of four in case of 16- and 32-channel PNPs, respectively. These clusters were also arranged equidistant along the longitudinal axis, with an inter-tetrode distance of approximately 50 µm. To achieve a dense array while facilitating microfabrication with a single metal layer in the flexMEA, electrical feedlines were downsized to a width and spacing of 2 µm. The conditions of a single metal layer and minimal width of the feedlines inevitably yielded wider flexMEAs in the case of the 32-channel PNPs compared to the 16-channel variants. Consequently, this results in a larger cross-section making the insertion of the 32-channel PNPs more challenging. The designs were iterated to reduce the footprint and the distance travelled by the carrier needle before the first electrode reached a target nerve fiber, enabling a reduction in the distance from the tip to the first electrode, from 59 – 995 µm to as little as 20 µm in 16- and 32-channel PNPs, respectively (Fig. 1C). The linear arrays offer straightforward spatial layouts that cover longer regions of the nerve, namely 462 µm and 539 µm, whereas the tetrode arrays cover 191 µm and 384 µm. However, the tetrode designs increase the local density of electrodes in a smaller region. Such denser electrode placement increases the likelihood of recording the action potentials of individual neurons or axonal fibers on multiple electrodes, which can be leveraged to isolate their respective activity through spike sorting ^20,29^.

The flexMEAs were bonded to custom-designed printed circuit boards (PCBs), which were then permanently glued to the carrier needle. This process involved manually aligning the flexMEA to the carrier needle under a microscope using a custom micromanipulator setup, which enabled the on-top bonding of the flexMEA on the carrier needle (Fig. 1D). Prior to alignment of the flexMEA, UV-curable glue was applied to the carrier needle. The adhesive’s wetting behavior on PaC created a smooth transition from the needle to the flexMEA (Fig. 1E; additional perspectives in Supplementary Fig. S1). The probe was further reinforced with a commercially available two-component epoxy to bind the carrier needle to the PCB. The total length of the assembly is approximately 5 cm, of which 2 cm are attributable to the flexMEA and the metallic carrier needle. The width of the carrier needle and flexMEA assembly is determined primarily by the width of the flexMEA, which is 220 µm and 435 µm for the 16- and 32-channel PNPs, respectively (Fig. 1F). The impedance magnitudes at 1 kHz of the bare Au electrodes ranged from 2.47 to 7.79 MΩ. To make the PNPs more suitable for neuronal recording, lower impedances are desirable ^30^. Therefore, a conductive coating based on roughened Au and PEDOT:PSS was electrodeposited in a post-processing step (Fig. 1F, inset).

The electrodeposition of Au was intended to increase the surface roughness of the electrode contacts and thereby enhance the adhesion between Au and PEDOT:PSS ^31^. Due to the increased surface area, the median impedance magnitude at 1 kHz was reduced by a factor of 4 after the Au electrodeposition. Subsequent electrodeposition of PEDOT:PSS further reduced the impedance magnitude, resulting in a total reduction of the median impedance magnitude at 1 kHz by a factor of 85, from an initial value of 5.45 MΩ down to 64 kΩ.

### Mechanical performance of the PNPs

To reach peripheral nerves for acute recordings in humans, PNPs need to be able to traverse through the skin and protective tissue layers, such as the epineurium. It therefore needs to be stable enough to withstand significant mechanical forces during the insertion and subsequent neural recording. To evaluate the mechanical and electrical stability of the PNPs, we conducted a series of tests using a commercially available AHS model ^32^ and repeated insertions into the exposed peripheral saphenous nerves from rats. We investigated three potential failure modes for the PNP. First, detachment of the flexMEA from the acupuncture needle. Second, loss of electrode viability after PNP insertion, and third, detachment of the PEDOT:PSS electrode coating which could impair recording quality and leave PEDOT:PSS residuals in the neural tissue.

To this end, we first confirmed that PNPs could be inserted into the AHS. Initial tests revealed that PaC-coated carrier needles underwent irreversible deformation because of deflections including buckling when penetrating the AHS, likely due to the slenderness of the needle and the reduced tip sharpness caused by the coating ^33,34^. While the deflection of the needle could have been mitigated by utilizing a larger carrier, the additional thickness could have increased the frictional force resulting in a higher load on the flexMEA and the adhesive ^35^. In general, the penetration and insertion of a multi-component object, such as the PNP, into inhomogeneous biological tissue constitute a complex modeling problem ^36^. Accordingly, it is challenging to determine an appropriate design for the PNP to ensure reliable insertion without causing damage to the PNP. To address this issue, we adopted a pre-holing approach, similar to those commonly used in experimental neural probe implantation in the central nervous system, to facilitate penetration of the dura mater ^37–39^. This approach involves piercing a hole in the skin with a cannula prior to inserting the PNP. We first pierced the AHS with cannulas of decreasing diameter (0.9 mm to 0.4 mm), thereby progressively increasing the mechanical resistance encountered during probe insertion (Fig. 2A), as small cannulas produce narrower channels through the skin. To assess the robustness of the adhesion between the flexMEA and needle, we inserted a batch of PNPs up to 10 times for each cannula diameter, resulting in a total of up to 40 insertions per probe. The largest cannula also included a plastic venous catheter which we used as a tunnel through the AHS to serve as the test case with the lowest possible mechanical resistance that could occur during probe insertion. Notably, the holes did not retain their initial size but underwent significant constriction due to the viscoelastic properties of the AHS, thus exerting additional shear forces on the PNP upon insertion.

**Figure 2.**
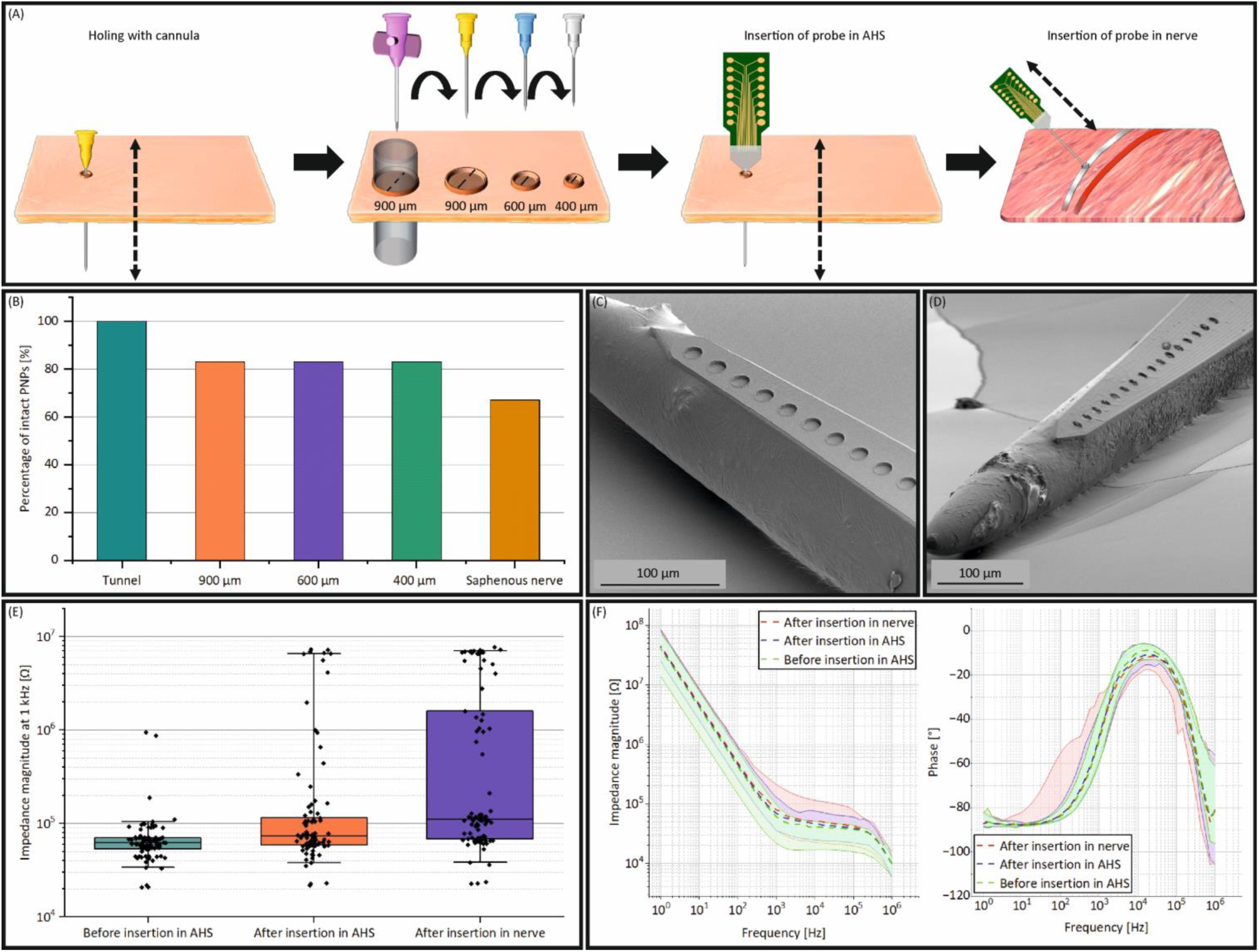
Mechanical and electrochemical performance of the PNP. **(A)** Schematic of mechanical tests of the PNPs, using cannulas to create differently sized holes in artificial human skin (AHS). The PNPs were then tested by inserting them through these holes. Notably, the holes contracted again after the initial penetration, requiring considerable force for the PNP to penetrate the AHS. Subsequently, the PNPs were also inserted into a rat saphenous nerve. **(B)** Bar plot depicting the percentage of mechanically intact PNPs after repeated insertion tests. The majority of probes stayed intact after all insertions. **(C)** SEM picture of a PNP before the mechanical stability test, exhibiting a seamless transition between the needle and flexMEA. **(D)** SEM picture of a PNP after the mechanical stability tests. The PNP structure was largely conserved with no visible signs of damage. **(E)** Box plot of impedance magnitudes at 1 kHz of each electrode before and after insertion trials. The boxes represent the inter-quartile range (IQR), the whiskers represent the 5%-trimmed ranges. **F)** Median and 5% trimmed ranges of electrical impedance of viable electrodes before and after insertions, illustrating the same qualitative impedance course.

We tested six PNP dummies and all insertions were performed manually by an experimenter, simulating microneurography needle insertion in the clinic. Each insertion was considered successful if no visible signs of mechanical alteration, such as delamination or rupture of the flexMEA, were observed on the PNP after insertion and extraction. From six dummy probes tested, all passed the initial test with the tunnel aid and only one PNP dummy showed signs of delamination for the first three hole sizes (Fig. 2B; exemplary delamination visible in Supplementary Fig. S2). After the skin penetration test, we proceeded with additional mechanical stress tests of the PNPs when entering the saphenous nerve of a euthanized rat. Here, the saphenous nerve was exposed while preserving the surrounding epineurium. The exposed nerve was then pierced manually under a surgical microscope with the same PNP dummies that were previously used in the AHS. Drying of the nerve and changes in tissue elasticity following euthanasia led to considerable mechanical forces on the device during perineum penetration (see Supplementary Video 1). Despite this, all but one PNP were successfully inserted and showed no visible signs of delamination or mechanical damage. This was confirmed by additional SEM imaging of representative PNP dummies before and after all mechanical testing stages, which consistently showed a seamless interface between the flexMEA and the carrier needle (Fig. 2C,D). These results corroborate that the flexMEA adhesion is sufficiently robust to withstand the high mechanical forces that occur during repeated insertions into skin models and nervous tissue.

These findings suggest that the PNPs are already mechanically robust but several fabrication- and handling-related factors can affect the mechanical stability of the PNPs. A key component is the adhesive and how it interacts with the (coated) needle and the flexMEA. This interaction may vary depending on the glue’s chemical composition. Specifically, different bonding mechanisms could increase adhesion, for example exploiting the properties of thermoplastic polymers for polymer-polymer bonding, thereby reducing the likelihood of detachment ^40,41^. The distribution of the glue, together with the alignment and thickness of the flexMEA, can also affect mechanical stability. As the flexMEA is an inherently protruding element, the cross-section of the PNP can change abruptly along the carrier needle at the tip of the flexMEA. This increases the likelihood of the flexMEA getting caught on the tissue during insertion. There are multiple reasons for such an abrupt transition: for example, the flexMEA could be non-negligibly thick or misaligned, or the glue could be unevenly distributed, failing to ensure a gradual transition. One potential strategy to mitigate the enlarged cross-section would be to adjust the shape of the flexMEA to the carrier needle by utilizing the thermoplastic property of PaC and a custom-made mold ^42^. Additionally, if the PNP is not properly aligned with the pre-hole in the skin during the insertion process, it will be necessary for the PNP to pierce the skin tissue, thereby, resulting in a higher mechanical load. Consequently, it would be advantageous to ensure an aligned insertion path, for instance by employing guiding pre-holing and insertion tools.

### Electrical performance

To assess the electrical stability of the PNPs, we used functional devices for penetrating the AHS using a 900 µm-wide pre-hole, followed by an insertion in the rat saphenous nerve. Microscopic inspections before and after the insertions confirmed that the PEDOT:PSS coating remained visually intact, with no apparent signs of delamination across the electrodes. The electrodes’ recessed location likely offered a distinct advantage, as it reduced the friction between the PEDOT:PSS-coated electrodes and the tissue during insertion, thereby decreasing the effective mechanical load (Supplementary Fig. S3).

To assess the electric viability of the electrodes an impedance magnitude of 1 MΩ at 1 kHz was pragmatically selected as an upper limit for an electrode to be considered functional. However, it should be noted that, depending on the electrode-axon coupling and the data acquisition system in use, signal quality may remain acceptable in some cases even with higher electrode-electrolyte impedances ^43,44^. Before insertion, the median of the impedance magnitude (|*Z*|) of the electrodes at 1kHz was 62.5 kΩ, with an inter-quartile range (IQR) of 53.7 – 70.7 kΩ. After testing, the impedance magnitude at 1kHz increased 1.2 fold (|*Z|* = 73 kΩ; IQR = [58.7 – 114.9 kΩ]) and 1.8 fold (|*Z|* = 110.7 kΩ; IQR = [68.5 – 1592.9 kΩ]) from the original value after insertion into the AHS and the nerve, respectively (Fig. 2E).

The electrodes exhibited a capacitive behavior at low frequencies (below 1kHz) and a resistive behavior at higher frequencies, as reported previously for PEDOT:PSS-coated electrodes (Fig. 2F) ^45^. Upon probe insertion and retrieval, an increase in impedance could be attributed to two potential factors. Firstly, a higher spread impedance was observed due to tissue residues (Supplementary Fig. S4). In this context, the spread impedance is the impedance resulting from tissue at the electrode-tissue interface ^46^. Secondly, there were impedance distortions akin to open-circuit impedances, which can be attributed to broken feedlines (Fig. 2F; individual electrode impedance spectra in Supplementary Fig. S5-S6). Notably, despite the impedance magnitude increases, 68% of the electrodes remained functionally viable (Fig. 2E, *N* = 90 electrodes), demonstrating that the electrical properties of the PNPs are largely preserved upon insertion into biologically relevant tissues.

It is also important to note that our results could even be an overestimate of the functional degradation of the electrodes because the impedance measures were done after both PNP insertion and extraction. Given that practical applications occur immediately after PNP insertion, the electrical performance of the PNP could be even higher under real-use conditions. In conclusion, these results demonstrate that the PNPs can withstand the mechanical stress encountered during insertion into biological tissue while largely maintaining their structural integrity and functional performance.

### *Ex vivo* validation

After confirming the PNPs mechanical and functional stability, we next assessed their performance for recording axonal nerve signals. We therefore conducted *ex vivo* nerve recordings, using a skin-nerve preparation from the rat saphenous nerve (Fig. 3A). This preparation is a well-established model for the extracellular recording of action potentials from peripheral sensory neurons, enabling controlled mechanical, thermal, or chemical stimulation of receptive fields in the skin to evoke neural activity in specific neural subtypes, such as mechanoreceptors and nociceptors. The saphenous nerve provides access to a broad population of afferent fibers, making it well-suited for validating neural probes *ex vivo* ^47,48^.

**Figure 3.**
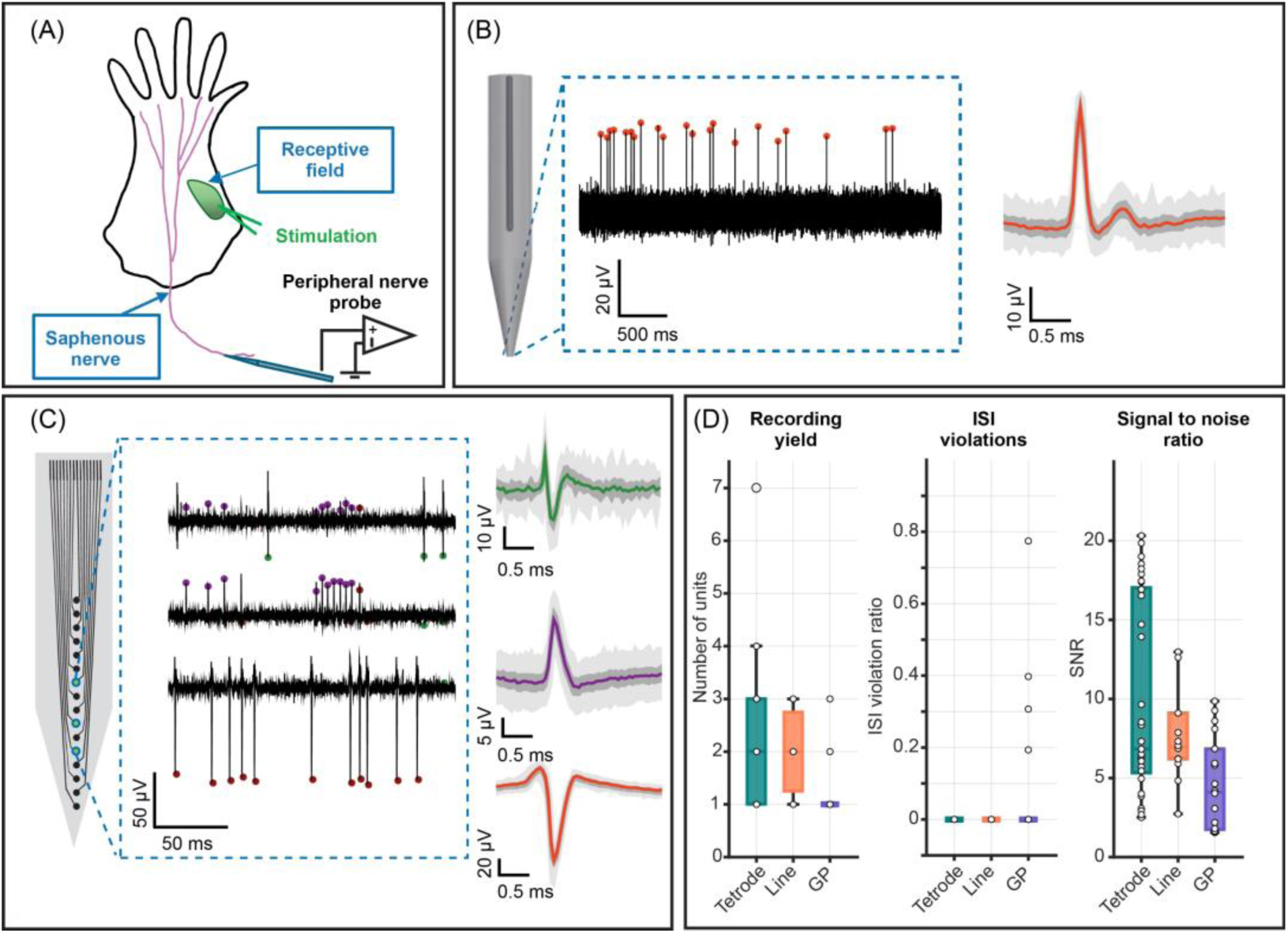
*Ex vivo* characterization of PNP configurations. **(A)** Schematic of the *ex vivo* skin–nerve preparation used for extracellular recordings. Mechanical stimulation of sensory neurons in the skin results in activation of axonal fibers in the saphenous nerve. Different probes (PNP_tetrode_, PNP_linear_, and GPs) were then used to record the spiking activity in individual nerve fibers. **(B)** Example of an electrophysiology recording from a GP. The black trace shows the high-pass filtered raw signal, traces on the right show the average action potential waveform for a single unit. Dark and light shading show the standard deviation and 99^th^ percentile across 500 waveforms, respectively. **(C)** Representative recording examples from a PNP_linear_. Traces show high-pass filtered raw signals with spiking activity from different axonal fibers. The traces on the right show average waveforms from multiple axonal fibers that were obtained with the PNP. Conventions as in B. **(D)** Comparison of the recording quality metrics across configurations. The number of detected units was highest with tetrode PNPs and lowest with GPs. ISI violation ratios were elevated in GP recordings, indicating poorer isolation of individual nerve fibers. Signal-to-noise ratio (SNR) was larger for the PNPs than for the GPs.

To assess PNP performance against an established standard for *ex vivo* nerve recordings, we compared 16-channel PNPs to glass micropipette suction electrodes (GP), which are widely used in skin-nerve preparations to study peripheral sensory neurons ^49^. The saphenous nerve was suspended inside a bath of synthetic interstitial fluid and recordings were made by sealing the pipette tip (∼170 µm opening diameter) to the nerve cross-section with mild negative pressure. For PNP recordings, the nerve was stabilized, and the probe was inserted from the proximal end using a piezo-driven micromanipulator. In both cases, receptive field mapping was then conducted by gently probing the skin surface with a glass rod to identify areas that evoked strong neural responses. This configuration enabled simultaneous detection of action potentials from multiple single axons and served as a benchmark to assess the performance of the PNP.

To evaluate the recording quality, we first used spike-sorting to isolate the activity of putative nerve fibers and manually curated the units, based on spike waveform shape, cluster separation in principal component space, and inter-spike interval (ISI) distributions. As part of this curation, we quantified the ISI violation rate, defined as the fraction of spikes violating the refractory period, as a practical indicator of cluster quality and false positive contamination.^50,51^ The curated dataset included 18 units from 14 GP recordings, 33 units from 15 PNP_tetrode_ recordings, and 14 units from 7 PNP_linear_ recordings. All three recording configurations (PNP_linear_, PNP_tetrode_, and GP) yielded action potentials from nerve fibers with high signal-to-noise ratios (SNRs) in response to mechanical stimulation (Fig. 3B,C). We observed variability in the spike waveforms, including positive, as well as bi- or triphasic spikes that likely originated from different nerve fibers. In human microneurography, negative triphasic waveforms are characteristic of unmyelinated C fibers and positive biphasic waveforms are indicative of myelinated A fibers ^16^. Different spike waveforms from an exemplary PNP_linear_ recording are shown in Fig. 3C, where spikes from at least three different nerve fibers were detected. Across all recordings, the unit yield also showed a small but significant difference between devices, with PNPs yielding a larger median amount of high-quality units per recording (Fig. 3D; Yield_tetrode_ = 2 units; IQR = [1 – 3 units]; Yield_linear_ = 2 units; IQR = [1.25 – 2.75 units]; Yield_GP_ = 1 unit; IQR = [1 – 1 unit]; Kruskal-Wallis test, *p* = 0.0234). Even after manual curation, some of the GP clusters also contained a larger degree of unphysiological ISI violations which are a sign of spikes from multiple fibers being combined in the same cluster ^52^ (Fig. 3D; ISI-Ratio_tetrode_ = 0; IQR = [0 – 0]; ISI-Ratio_linear_ = 0; IQR = [0 – 0]; ISI-Ratio_GP_ = 0; IQR = [0 – 0.17]; Kruskal-Wallis test, *p* = 0.0042). Lastly, we also assessed the signal quality across recording configurations by comparing the SNR of curated units. In line with our previous results, the units from PNP recordings showed significantly higher SNRs than units from GP recordings, underscoring the utility of the PNPs for high-quality extracellular nerve recordings (Fig. 3D; SNR_tetrode_ = 6.83; IQR = [5.33 – 17.04]; SNR_linear_ = 6.96; IQR = [6.2 – 9.1]; SNR_GP_ = 4.11; IQR = [1.74 – 6.85]; Kruskal-Wallis test, *p* = 0.0043).

Together, these results clearly demonstrate that the PNPs are very well-suited for extracellular recordings of axonal action potentials in peripheral nerves and may offer several advantages over GPs. GPs pose several practical and technical drawbacks, including mechanical fragility, limited spatial coverage, and a general lack of scalability to multi-site recordings. In contrast, PNPs provide mechanically robust, scalable, and geometrically versatile alternatives that allow for greater flexibility in targeting peripheral nerves. By arranging multiple recording sites in defined linear or tetrode configurations, the PNPs also enable more structured sampling across nerve fascicles and increase the likelihood of isolating single-unit activity from individual axons.

### *In vivo* validation in a rat model

In order to assess the performance of PNPs under physiological conditions, we conducted acute *in vivo* experiments targeting the saphenous nerve in anesthetized rats (Fig. 4A). For these experiments, we used a 32-channel version of the PNP (Fig. 4B) with a similar electrode arrangement as the 16-channel PNP_tetrode_ used *ex vivo*. This layout increased electrode density within the nerve while also extending the sensing area and thereby improving the likelihood of capturing axonal activity across a broader region of the nerve.

**Figure 4.**
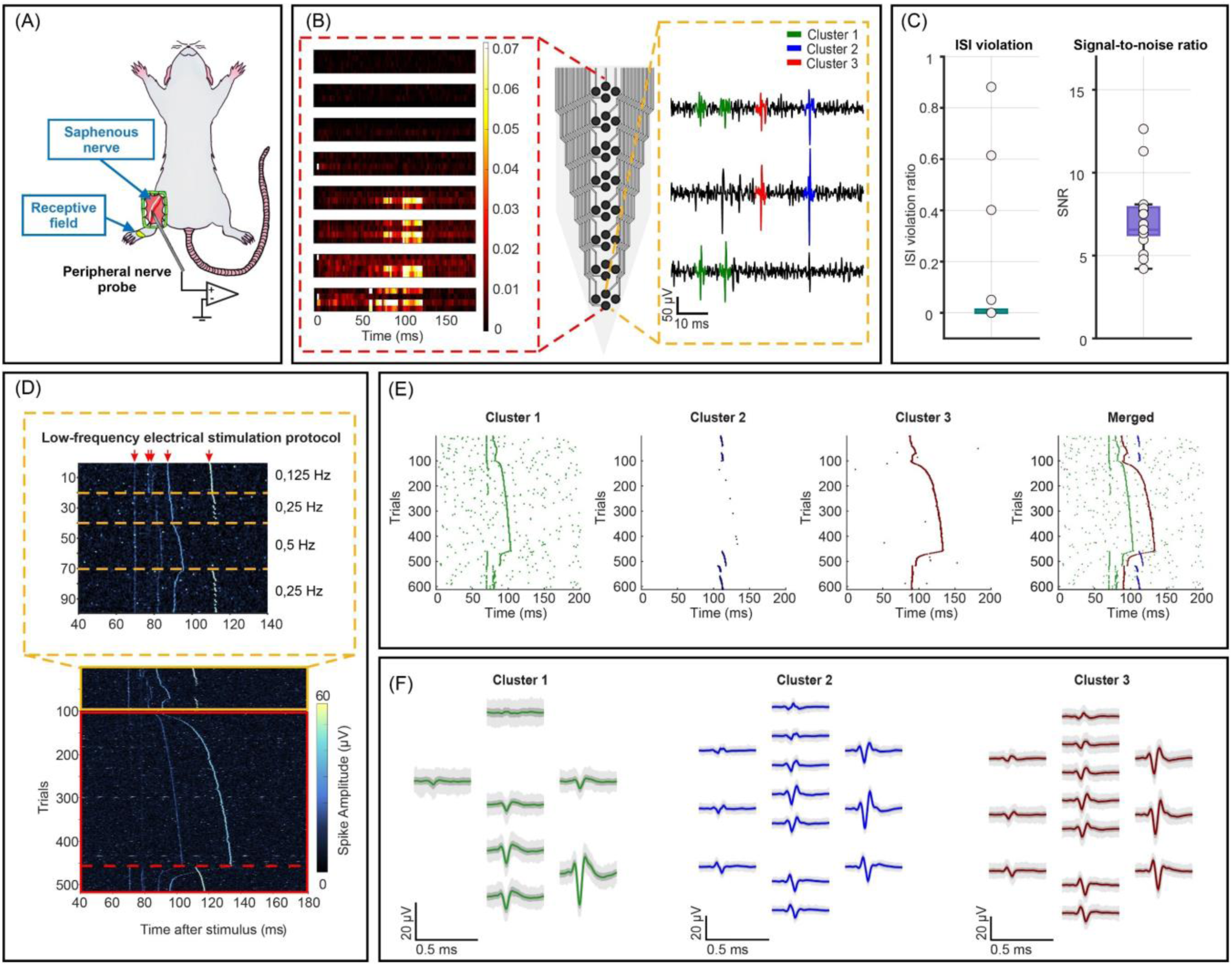
*In vivo* characterization of peripheral nerve probes. **(A)** Scheme of *in vivo* experiment of recording activity of peripheral nerves. **(B)** Left: Spatial distribution of electrically-evoked and averaged neural activity across channels. Center: Layout of the 32-channel PNP_tetrode_. Right: Example band-pass-filtered traces after common average re-referencing. **(C)** ISI violation ratios and SNR across all spike-sorted clusters. **(D)** Trial-by-trial latency variation heatmap during electrical stimulation showing multiple candidate C-nociceptor and C-non-nociceptor fibers on individual recordings *in vivo*. Yellow dashed rectangle (top) shows results from the low-frequency stimulation, the red rectangle (bottom) from the high frequency stimulation protocol. **(E)** Functional separation of fibers, based on spike sorting with MountainSort 5. Right panel shows merged responses from all three clusters. **(F)** Corresponding waveforms of the identified fibers in E).

Following the PNP insertion into the nerve, we used mechanical stimulation to identify the receptive fields of sensory neurons. We then electrically stimulated the skin in the mechanical receptive field, from which we could record high-quality axonal signals in the nerve, characterized by consistent stimulus responses and a good SNR (Fig. 4C). Electrical stimulation enables precise measures of stimulus-to-spike latency and latency change of signal conduction, which can be used to distinguish different fiber subtypes, such as Aδ- and C-fiber nociceptors ^7,12^. We used low- and high-frequency stimulation protocols to induce different latency changes and functionally characterize the recorded nerve fibers. A clear time-locked response to stimulation was visible across multiple channels, particularly on electrodes near the probe tip that were positioned deeper in the nerve. In addition, distinct spike waveforms were detected across channels, consistent with multiple fibers being recorded simultaneously (Fig. 4B, Supplementary Figure S7). The low-frequency stimulation provided a baseline measure of excitability, while the stepped high-frequency protocol was used to probe for potential activity-dependent slowing. Both low and high-frequency stimulation protocols have been shown to distinguish different classes of C-fiber nociceptors in pigs and humans ^15,53^. Trial-by-trial latency analysis revealed a progressive change in latency with repeated stimulation for some fibers (Fig. 4D, Supplementary Fig. S7B), consistent with the recruitment of distinct fiber types, including C-fiber nociceptors displaying activity-dependent slowing.

To evaluate the recording quality *in vivo*, we performed spike sorting using the SpikeInterface framework ^23^. Electrical stimulation artifacts were removed, and the raw neural data were band-pass filtered and re-referenced to the common median over all channels. Multiple spike sorting algorithms were applied to detect and classify neuronal units, including Kilosort 4, Tridesclous 2, Spiking circus 2, and MountainSort 5. All sorters yielded a relatively high number of clusters, consistent with the complexity of peripheral nerve recordings *in vivo*, which often involve overlapping waveforms and high background activity. However, the total number of clusters differed substantially across sorters (MountainSort 5: 18, Kilosort 4: 26, TrideClous 2: 44, Spiking circus 2: 32), suggesting high variability in spike sorting performance. We assessed the quality of the detected clusters by examining their latency stability to electrical stimulation across trials. MountainSort 5 yielded the greatest proportion of physiologically interpretable units, based on their consistent time-locked spike latency in response to electrical stimulation, waveform quality, and cross-fiber response separation (Fig.4). The best three clusters from other sorters are shown in Supplementary Figure S8. Further manual curation was based on waveform shape, ISI distribution, spike count, and trial-by-trial latency stability, using the Phy2 interface ^54^.

Units from *in vivo* recordings had slightly higher median ISI violation ratios compared to *ex vivo* recordings (Fig. 4C; ISI-Ratio*in vivo* = 0; IQR = [0 – 0.04]; *n*_clusters_ = 17), likely due to the more challenging recording environment. Additionally, signal quality was quantified by calculating the SNR of curated units. Consistent with our previous findings, units from PNP recordings *in vivo* demonstrated SNRs comparable to those from *ex vivo* recordings, highlighting the robustness of PNPs for extracellular nerve recordings under more complex experimental conditions, where factors such as tissue motion and physiological noise can influence signal quality. (Fig. 4C; SNR = 6.56; IQR = [6.33 – 7.85]; *n*_clusters_ = 17).

In a representative recording, we identified three well-isolated single units that exhibited reliable responses to electrical stimulation (Fig. 4E), as well as nine additional units displaying spontaneous firing activity (Supplementary Fig. S9). Functional units exhibited ISI violation rates below 1 (range: 0–0.4), indicating clean, single-fiber isolation. Waveform similarity between some clusters also highlighted that spike shape alone is insufficient to distinguish some units due to the spatial proximity and size similarity of axons. This phenomenon is illustrated in Figure 4E-F. Cluster 1 shows overlapping waveforms but with different latencies to electrical stimulation, suggesting a merged unit that likely contained the activity of at least two axons. This unit can be further split by their latency change. In contrast, clusters 2 and 3 demonstrate clearly separated waveform features and highly consistent response latencies, supporting their classification as individual fibers. The present findings demonstrate the technical feasibility of isolating single-unit activity from peripheral nerves *in vivo* using PNPs. Nonetheless, the existing spike sorting algorithms are constrained in their ability to accurately separate closely spaced axonal signals with similar waveforms and localization across the same probe contacts.

## Conclusion and Outlook

In this work, we developed a mechanically and electrically stable multi-contact neural probe for acute recordings in peripheral nerves. The PNPs are based on flexible MEAs which are glued to acupuncture needles in a custom-built alignment setup. Their mechanical stability was demonstrated by repeated insertions through the AHS model and into saphenous nerves of rats while their electrical stability was confirmed by additional insertion experiments and electrical measurements.

Our findings demonstrate that PNPs enable stable, multi-site recording of evoked neural activity *ex vivo* and *in vivo*, with sensing capabilities comparable to conventional multisite silicon probes. Moreover, their spatial electrode configurations and enhanced mechanical stability offer key advantages for long-duration acute recordings and functional mapping of peripheral nerve fibers. These features, along with the use of biocompatible materials make the device well-suited for experimental and clinical applications requiring minimal invasiveness and high mechanical stability, with the potential for translation to acute clinical diagnostics in humans.

Future research will focus on enhancing the mechanical and electrical stability of the PNPs, and increasing their spatial resolution through denser arrays and reduced electrode diameters. Scaling electrode dimensions to match those of typical mammalian nerve fibers, such as unmyelinated C fibers (0.2–1.5 µm in diameter) and thinly myelinated Aδ fibers (1-6 µm) ^7,11^, is expected to further enhance their ability to effectively capture the activity of individual nerve fibers. Furthermore, applying PNPs in larger animal models and diverse disease contexts will be essential to evaluate their robustness, versatility, and translational potential towards human applications. Ultimately, we envision PNPs as a valuable substitute for conventional microneurography needles for studying pain and neurological disorders, enabling advances in peripheral recordings and improving the physiological characterization and diagnosis of peripheral nerve pathologies.

## Materials and Methods

### PNP assembly

#### Fabrication of flexMEA

The flexMEAs were fabricated in cleanroom environments^55^. First, a layer of Parylene-C, 3–6 µm thick, was deposited on 4-inch silicon wafers in the deposition tool PDS 2010 Labcoter 2 (Specialty Coating Systems, USA).

Next, the metal layer was formed using a lift-off process. Initially, the wafers were conditioned on a hotplate set to 150 °C for 5 minutes. Subsequently, 3.5 ml of the negative photoresist AZ LNR-003 (MicroChemicals GmbH, Germany) was spin-coated onto the wafers. The wafers were spun at a speed of 4000 rpm for 45 seconds with an acceleration of 500 rpm/s. A soft bake step of two minutes was then carried out on a hotplate set to 120 °C. Light exposure to implement the patterns was carried out either using the maskless direct writer MLA 150 (Heidelberg Instruments Mikrotechnik GmbH, Germany) or the mask aligner MA/BA 8 (SUSS MicroTec SE, Germany). For the maskless direct writer, a dose of 320 mJ/cm² and a defocus setting of 2 were used. A critical dimension bias of 800 nm was applied to the exposure pattern. For the mask aligner, doses of either 120 or 130 mJ/cm² were applied. A post-exposure bake was then carried out on a hotplate for 90 seconds. For photoresist exposed in the mask aligner, the temperature was set to 110 °C. For the photoresist exposed in the maskless direct writer, the temperature was set to 100 °C. Development was carried out in the developer solution AZ 326 MIF (MicroChemicals GmbH, Germany). In the case of exposure at the mask aligner, the development time was 60 seconds. In the case of exposure at the maskless direct writer, the development time was 90 seconds.

Metals were deposited by electron beam-assisted evaporation in a physical vapour deposition process. The following layers were deposited from bottom to top: 20 nm of titanium, 100 nm of gold, and 10 nm of titanium. The deposition rates were set to 0.1 nm/s for titanium and 0.5 nm/s for gold. The lift-off process was completed by removing the photoresist together with the excess metal. For this purpose, either acetone or TechniStrip NI555 (MicroChemicals GmbH, Germany) was used at an elevated temperature.

After the lift-off process, a second PaC layer was deposited in a manner analogous to the first layer. To obtain the shape of the flexMEAs and create openings for the electrodes and bond pads in the passivation layer, a dry etching process was performed. To achieve this, an etch mask consisting of the positive photoresist AZ 12XT (MicroChemicals GmbH, Germany) was created. The wafers were first conditioned on a hotplate set to 150 °C for 5 minutes. Then, 5 ml of photoresist was applied on each wafer. The wafers were spun at 1000 rpm for 3 minutes. The acceleration was set to 200 rpm/s. A soft bake was then carried out on a hotplate set to 110 °C. The soft bake duration was either 4 or 5 minutes. Light exposure for patterning was carried out in either the aforementioned mask aligner or maskless direct writer. For exposure using the mask aligner, a dose of 180 mJ/cm² was applied. For the maskless direct writer, a dose of 350 mJ/cm² was applied, along with a defocus setting of 2. A critical dimension bias of -800 nm was applied to the exposure pattern. The post-exposure bake was carried out on a hotplate set to 90 °C for 60 seconds. Development was carried out in AZ 326 MIF for 2 minutes.

Dry etching was performed using an inductively coupled plasma reactive ion etching tool (Oxford Instruments Plasma Technology, UK). The power of the plasma generator and the DC bias generator were set to 500 W and 50 W, respectively. Oxygen (O₂) and carbon tetrafluoride (CF₄) were used with flow rates set to 36 and 4 sccm, respectively. The chamber pressure was set to 0.7 Pa and the backside cooling temperature of the wafer was set to 10 °C. The etching times were calculated linearly with an assumed etch rate of 800 nm/min. The etch mask was then removed using AZ 100 remover (MicroChemicals GmbH, Germany).

#### Packaging

Following microfabrication, the flexMEAs (MEAs) were removed from the substrate wafers. Water was used to aid this step. The flexMEAs were then manually soldered onto PCBs in a process analogous to the flip-chip bonding process. The metal composition of the solder paste was 42% tin and 58% bismuth. After soldering the flexMEA to the PCB, the corresponding connector was soldered to the PCB. For probes with 16 electrodes, the pins of DIL connectors were soldered onto the PCB. For probes with 32 electrodes, SAMTEC MOLC-110-01-S-Q connectors were soldered onto the PCB.

#### On-top bonding

The acupuncture needles utilized as carrier needles were type J acupuncture needles with a shank diameter of 120 µm and a length of 3 cm (SEIRIN Corporation, Japan). A custom-built alignment setup was used to bond the flexMEAs onto the carrier needles. It consisted of a pair of self-closing tweezers to hold the carrier needle and a micromanipulator with an arm to which a flexMEA could be mounted. Firstly, the medical-grade, UV-curable glue Vitralit 7311 FO (Panacol-Elosol GmbH, Germany) was applied to the carrier needle. The flexMEA was then aligned with the carrier needle using a microscope for visual inspection. After alignment, the assembly was exposed to UV light to cure the glue and fix the flexMEA to the carrier needle. In a second step, the carrier needle handle was affixed to the PCB using commercially available two-component epoxy glue (UHU GmbH & Co. KG, Germany).

#### Electrode coating

In the case of electrodes with an additional, electrodeposited gold layer, the PNPs were immersed in chloroauric acid. The initial concentration of the solution was 100 mM. A three-electrode setup with a platinum wire as the counter electrode, an Ag/AgCl pseudo-reference electrode as the reference electrode, and the electrodes of the PNPs as the working electrode was set up. A potential of -0.2 V was applied for 20 seconds in a chronoamperometry process, carried out using the potentiostat CHI1030B (CH Instruments, Inc., USA).

Prior to PEDOT:PSS electrodeposition on probes without an additional gold-electrodeposited layer, the electrodes underwent 11 cycles of cyclic voltammetry in 1x phosphate-buffered saline (PBS) within a voltage range of -0.6 V to 0.9 V, at a sweep rate of 0.1 V/s. The process was carried out using the potentiostat VSP-300 (BioLogic, France). After this, the PNPs were treated with an oxygen plasma for 3 minutes. 80 W of power was used to generate the plasma and the chamber pressure was set to 0.8 mbar. This was done using the plasma oven Pico (Diener electronic GmbH & Co. KG, Germany). PEDOT:PSS deposition was conducted in an EDOT:PSS solution containing 3,4-Etyhlenedioxythopene (CAS: 126213-50-1; Sigma-Aldrich Chemie GmbH, Germany) and poly(4-styrenesulfonic acid) solution (CAS: 28210-41-5; Sigma-Aldrich Chemie GmbH, Germany). It was deposited in a chronoamperometry process with the same, above-mentioned three-electrode setup that was used for the electrodeposition of gold. A potential of 1 V was applied to the electrodes immersed in the EDOT:PSS solution for 20 s.

### Characterization

#### Electrochemical

Electrochemical characterization was carried out by electrochemical impedance spectroscopy in the same three-electrode setup that was used for electrode coating. The potentiostat VSP-300 (BioLogic, France) was used here. The impedance was measured in the frequency range from 1 Hz to 1 MHz in 1x PBS. The applied voltage amplitude was 10 mV.

#### Mechanical stability

The AHS model Double Layer Skin with the thin dermis option (LifeLike BioTissue, Canada) was fixed on a beaker filled with water. The skin model was kept moist and wetted every 25 to 30 min with water during the experiments.

### *Ex vivo* validation

#### Ethical statement

Use of tissues in this work has been approved by the state animal ethics committee, the Landesumweltamt für Natur, Umwelt und Verbraucherschutz Nordrhein-Westfalen, Recklinghausen, Germany, under approval names: IBI-3_OE_Ratte and IBI3-OE Ratten2_VRM. It has been conducted according to local animal protection regulations and is reported according to the ARRIVE guidelines.

#### Skin-Nerve Preparation

The experimental procedures were performed on adult Wistar rats. Anesthesia was induced using 5% isoflurane for a minimum of five minutes and maintained until the absence of the paw withdrawal reflex. Following anesthesia, animals were euthanized via decapitation. The bodies were then transferred to a designated surgical area.

The inner sides of both hind limbs were shaved from the ankle to the groin to improve access. One hind limb was fixed on an angular stand using double-sided tape. Perpendicular incisions were made near the base of each toe, extended proximally along both sides, and the top layer of the skin was elevated with an incision made near the bone to preserve nerve branching and terminals. Additional incisions along the lateral side of the paw released the dorsal skin.

To expose the saphenous nerve, an incision was made in the femoral region to identify the femoral artery and associated nerve. Lateral and distal incisions were extended to connect with previous cuts. Connective tissue between the skin and muscle was removed, and a longitudinal incision was made along the nerve’s path to free it down to its junction with the skin. The preparation was frequently rinsed with synthetic interstitial fluid (SIF). The fully dissected skin-nerve complex was immersed in carbogenized (95% O₂, 5% CO₂) SIF with the following composition (in mM): NaCl 107.7, KCl 3.48, NaHCO₃ 26.2, NaH₂PO₄ 1.67, Na-gluconate 9.64, glucose 5.55, sucrose 7.6, CaCl₂·2H₂O 1.53, MgSO₄·7H₂O 0.69. All the experiments were carried out at room temperature.

#### Recording Setup

The skin-nerve tissue was transferred to a recording chamber filled with carbogenized SIF and maintained under continuous perfusion via a peristaltic pump. The chamber bottom was precoated with Vaseline to secure the tissue. The nerve was carefully desheathed prior to electrode application.

- *Glass Micropipette Approach (GP)*: The nerve was left floating, and a glass micropipette was guided to the nerve tip using a magnetically mounted rotational holder. Negative pressure was applied to seal the pipette tip onto the nerve cross-section. Receptive fields were located by gently probing the skin surface with a glass rod.
- *Peripheral Nerve Probe (PNP) Approach*: The nerve was pinned at both ends—distally at the tip and proximally via muscle insertion. The PNP was inserted into the nerve using a piezo-driven micromanipulator (Sensapex, Finland). Receptive field searches were conducted as above. Both electrode configurations were tested: tetrode and linear.

Recordings were acquired using an in-house developed BioMAS acquisition system, consisting of a BioMAS amplifier, 16-channel MEA16.II.1 headstage, NI USB-6255 ADC, and custom software. Signals were sampled at 20 kHz and visualized in BioMAS Viewer. Working electrodes exhibited impedance magnitudes < 1 MΩ at 1kHz (ranging from 28.5 kΩ to 232 kΩ, *n* = 117 electrodes from 8 PNPs).

### *In vivo* validation

#### Ethical statement

All experimental procedures complied with the UK Animals (Scientific Procedures) Act 1986 and received approval from the University of Bristol Animal Welfare and Ethical Review Body. Rats were housed in groups under a 12-hour light/dark cycle, with food and water provided ad libitum. Saphenous nerve recordings were obtained from two adult male Wistar rats under terminal anaesthesia.

#### Surgery

The methods were similar to those described previously in detail for silicon based multisite probe peripheral nerve recordings ^4^. Anaesthesia was induced with 5% isoflurane until the loss of paw withdrawal reflex was observed. Subsequently, the animal was positioned on the heating mat, and a rectal temperature detector was inserted. The rectal temperature then was maintained to be at 37° C. The hind limb was then fixed to the surgery surface using double-sided tape. Then the area was prepared by shaving the fur from the groin crease to the ankle, followed by thorough cleaning and disinfection. The skin incision was made distally, along the femoral part of the leg, to access the saphenous nerve. The connective tissue between the skin and the muscles was then removed, and the skin ends were connected to the metal frame using interrupted sutures to create a “pool” that was filled with warm (37-40° C) PBS to prevent desiccation. Then, under the microscope, a small segment of the nerve was blunt dissected free from the surrounding tissues and blood vessels. A small incision was then made along the epineurium using a dull cannula tip to allow better access to nerve fibers.

### Recording setup

Neural activity was recorded using 32-channel PNPs arranged in a tetrode electrode configuration. Data acquisition was performed with the OpenEphys system, paired with a 32-channel headstage (Intan, RHD 2132). Each channel was sampled at 30 kHz with a gain of 192. The probe and headstage were linked *via* a Neuronexus headstage adaptor (#A32-OM32). The recorded signals were processed using the OpenEphys software (https://open-ephys.org/).

To ensure precise placement, the probe was mounted on a hydraulic micromanipulator (Narishige) and inserted into the nerve at a shallow angle. The nerve was carefully stabilized using fine forceps gripping the connective tissue sheath, minimizing the risk of neural damage. The probe was then gradually advanced into the exposed nerve while data visualization was provided by the OpenEphys graphical user interface. A signal chain was configured to apply a bandpass filter (300–3000 Hz) and common average referencing. Successful insertions were characterized by the immediate appearance of action potentials across multiple channels.

Once neural units were observed, a soft paintbrush was gently brushed against the dorsal surface of the foot to identify the skin region innervated by the fascicles of the saphenous nerve being recorded from. When receptive fields were identified, responses to electrical stimulation were assessed using bipolar transcutaneous needle electrodes connected to a constant current stimulator (DS4, Digitimer, UK). The stimulator was controlled via TTL pulses generated by an Arduino Uno running custom scripts to vary stimulation intervals. Electrical pulses of 3.2 mA and 0.5 ms duration were applied to activate axon terminals within the identified receptive fields. To induce latency changes, we used a low-frequency stimulation protocol that included 20 pulses at 0.125 Hz, 20 pulses at 0.25 Hz, 30 pulses at 0.5 Hz, and an additional 20 pulses at 0.25 Hz. After a two-minute rest period, a high-frequency protocol was applied, consisting of three minutes of continuous stimulation at 2 Hz, followed by 60 pulses at 0.25 Hz. ^4^

At the end of the experiment, the animal was euthanized by cervical dislocation under deep anaesthesia.

#### Data and Statistical Analysis

The raw data of the *ex vivo* experiments were filtered with a high-pass filter with a cut-off frequency of 300 Hz. The data of the *in vivo* experiments were band-pass filtered to a bandwidth from 300 Hz to 3000 Hz, followed by common average references. Spike sorting was performed using MountainSort5 *via* Spike Interface framework^23^, which uses an iso-split clustering algorithm based on unimodal distribution detection. Manual inspection and curation of sorted units were conducted using Phy2^54^ to ensure accuracy and eliminate false positives. Clusters were evaluated based on their waveform features and principal components, and for *in vivo* data, also based on the latency change for each cluster. We excluded all units exhibiting more than 1 ISI violations or signal-to-noise ratios (SNR) near 1, indicative of low-quality or spurious units, from further analysis.

Three probes (GP, PNP_linear_, and PNP_tetrode_) were evaluated using: number of detected units, signal-to-noise ratio (SNR), and inter-spike-interval (ISI) violation ratio. The one-sided Mann–Whitney U test and non-parametric statistical tests were applied throughout.

Schematics were generated using Blender v.4.2 (Blender Foundation, Netherlands) and CorelDRAW (Graphics Suite 2025, Corel Corporation, Canada). OriginPro (OriginLab Corporation, USA) and self-written scripts in MATLAB vR2024a (The MathWorks, Inc., USA) were used for data analysis.

## Supporting information

PNP_Supplementary information

## Acknowledgments

The authors thank the Helmholtz Nano Facility (HNF) at *Forschungszentrum* Jülich for facilitating the microfabrication of the devices. The authors thank Prof. A. Offenhäusser for infrastructural and scientific support. The authors thank B. Breuer for organizational support, E. Brauweiler-Reuters for carrying out SEM, and M. Prömpers, M. Banzet, and R. Stockmann for technical support at the HNF. This work was supported by the Helmholtz Association (VH-NG-1611), the Volkswagen Foundation (A.z.9C024), and the Exploratory Research Space of the RWTH Aachen University (OPSF640).

## Contribution

B.N., S.M., and V.R.M. conceived the development of the PNP probes and planned the study. E.Y and V.R.M. fabricated the devices. E.Y. and S.D. carried out the electrochemical and mechanical characterization of the devices with the support of S.M. and V.R.M. S.D. conducted *ex vivo* experiments with the support of S.K., B.N., S.M., and V.R.M. J.D. conducted the *in vivo* experiments on rats with the support of E.Y., S.D., A.E.P., B.N., and V.R.M. S.D. and E.Y collected and processed data and figures with the support of A.J., S.M., and V.R.M. S.D., E.Y., S.M., and V.R.M. wrote the initial draft of the manuscript. All authors reviewed and edited the manuscript. B.N., S.M., and V.R.M. supervised the project.

