## Supplementary material for "A mechanically stable neural probe for percutaneous high-resolution, multi-channel recordings in peripheral nerves": PNP_Supplementary information

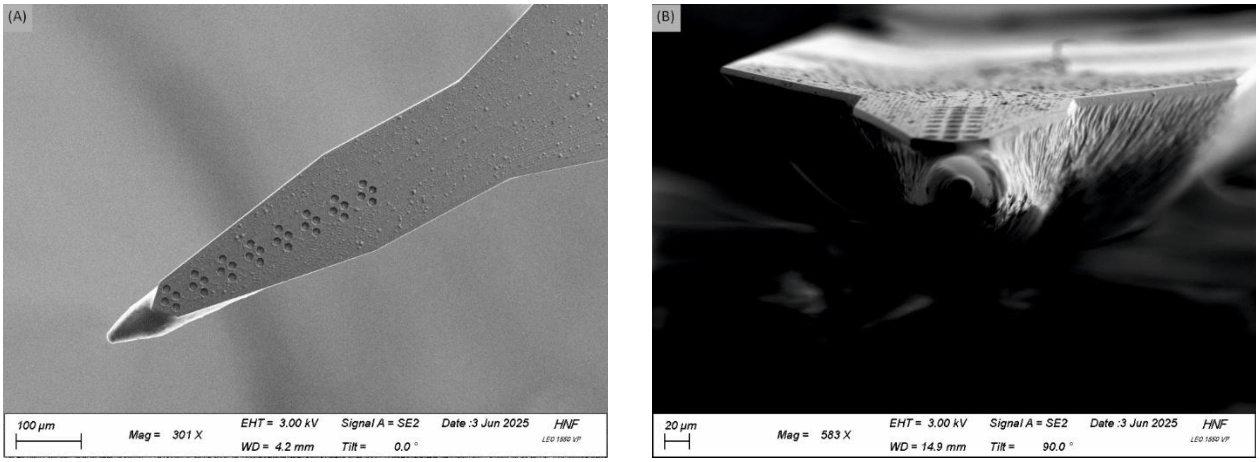


Figure S1: SEM pictures of 32-channel PNP. A) Top-Down view. B) Frontal view depicting the contact angle between the glue and the flexMEA.

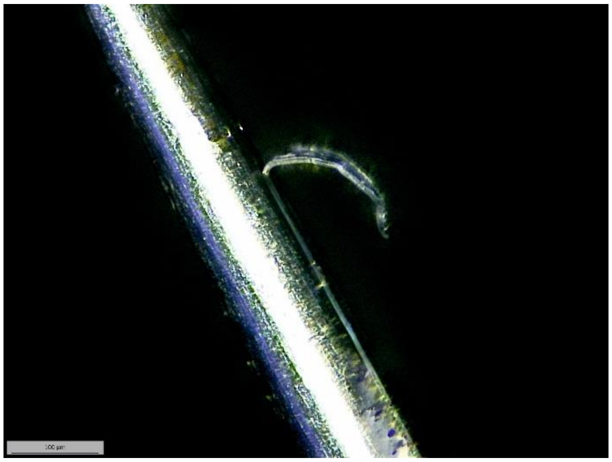

Figure S2: Microscopy image of PNP with detached flexMEA after insertion attempt.

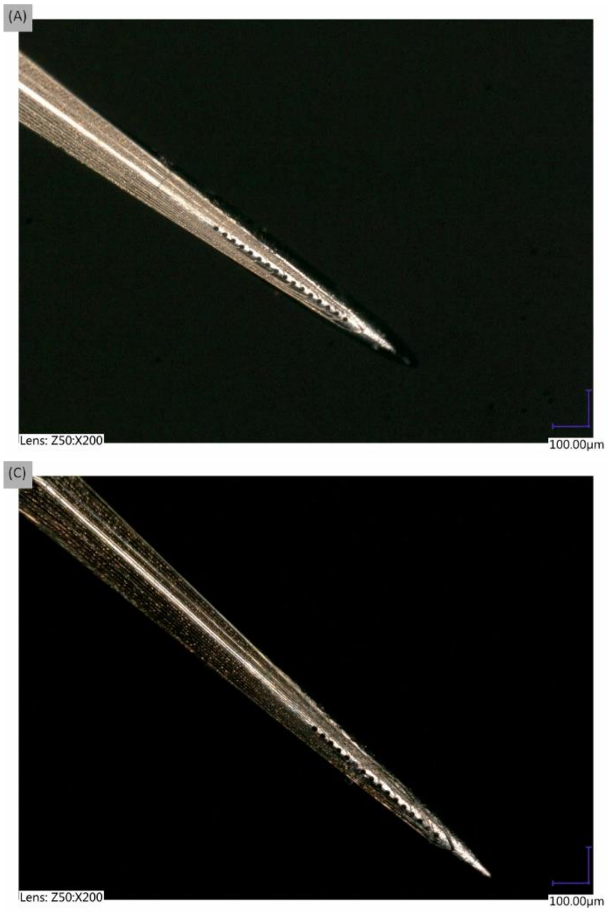


Figure S3: Microscopy images of an exemplary PNP with PEDOT-PSS electrode-coating. A) Before any insertion. B) After insertion through in AS with a 900 µm-wide hole. C) and D) After insertion in saphenous nerve. Some of the electrodes are covered by tissue after the insertion into the saphenous nerve.

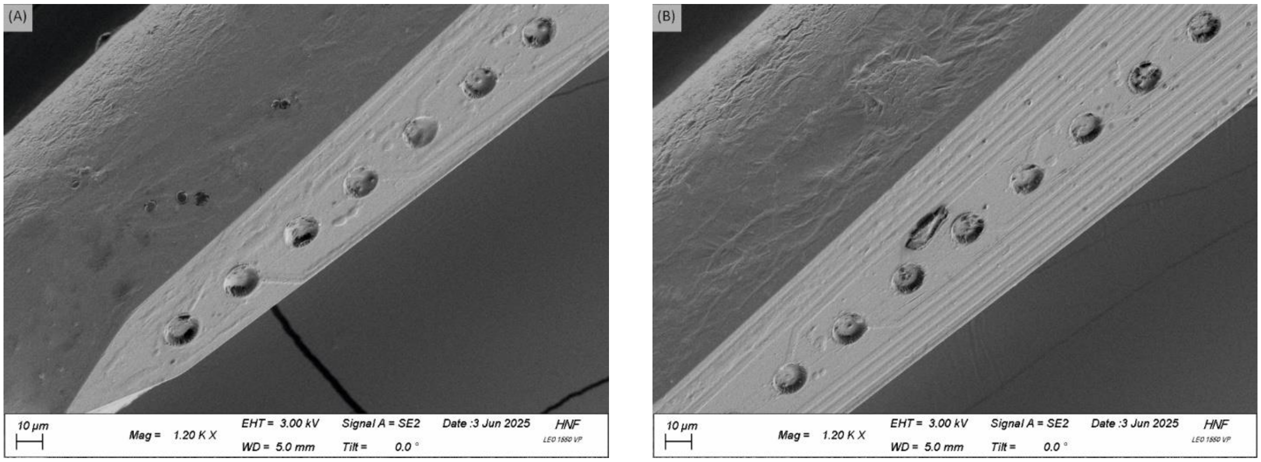


Figure S4: SEM pictures of 16-channel PNP after insertion into saphenous nerve. Electrode openings seem to be filled with material, possibly biological tissue from the insertion test. A) Tip area of flexMEA. B) Section after tip area of flexMEA.

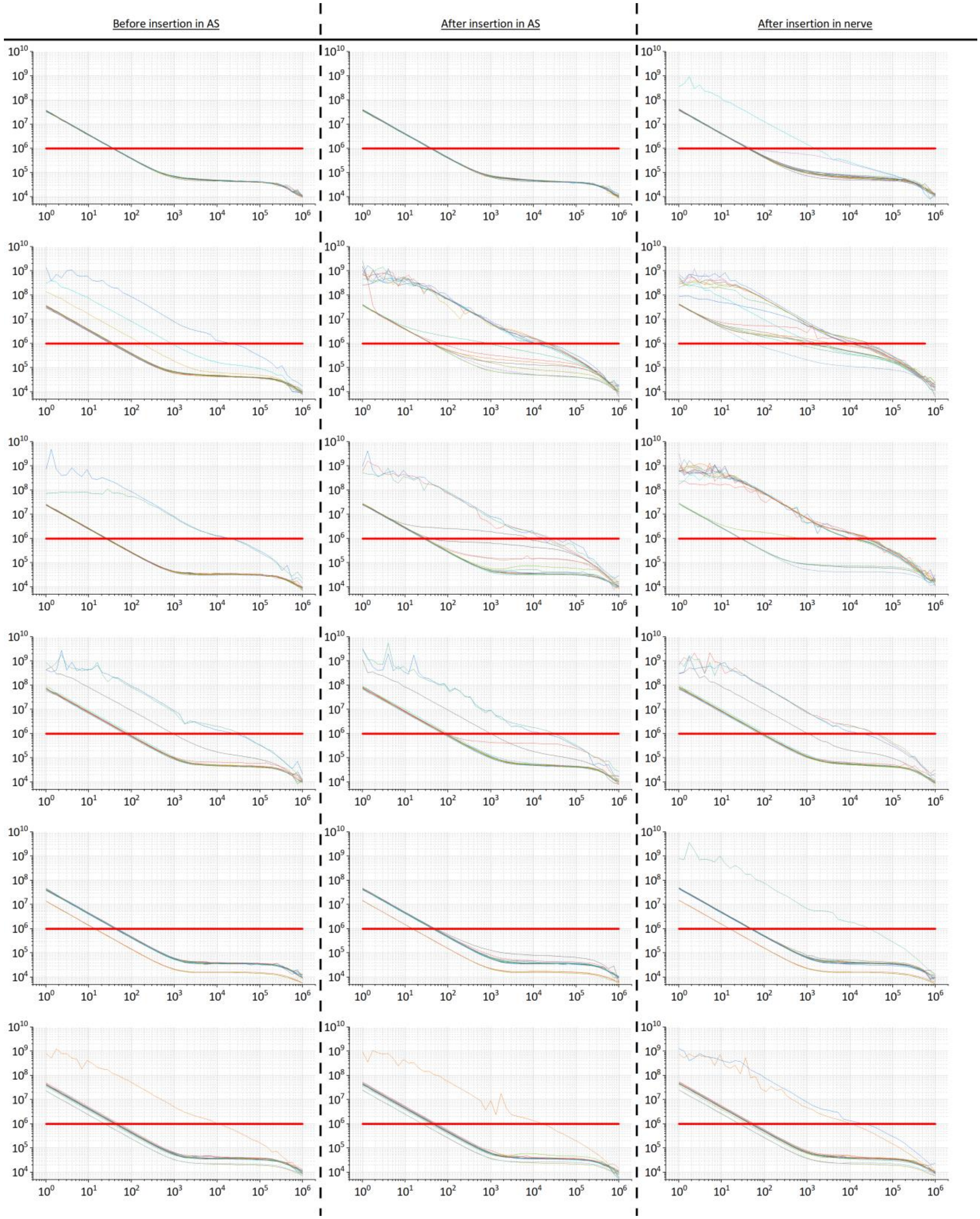


Figure S5: Impedance magnitudes of electrodes from probes used for electrochemical stability tests. X-axes represent frequency [Hz] and y-axes represent magnitude [Ω]. Red, horizontal line at 1 MΩ marks viability threshold. Each of the 6 rows belongs to one individual probe. The order of probes is the same as in Figure S6.

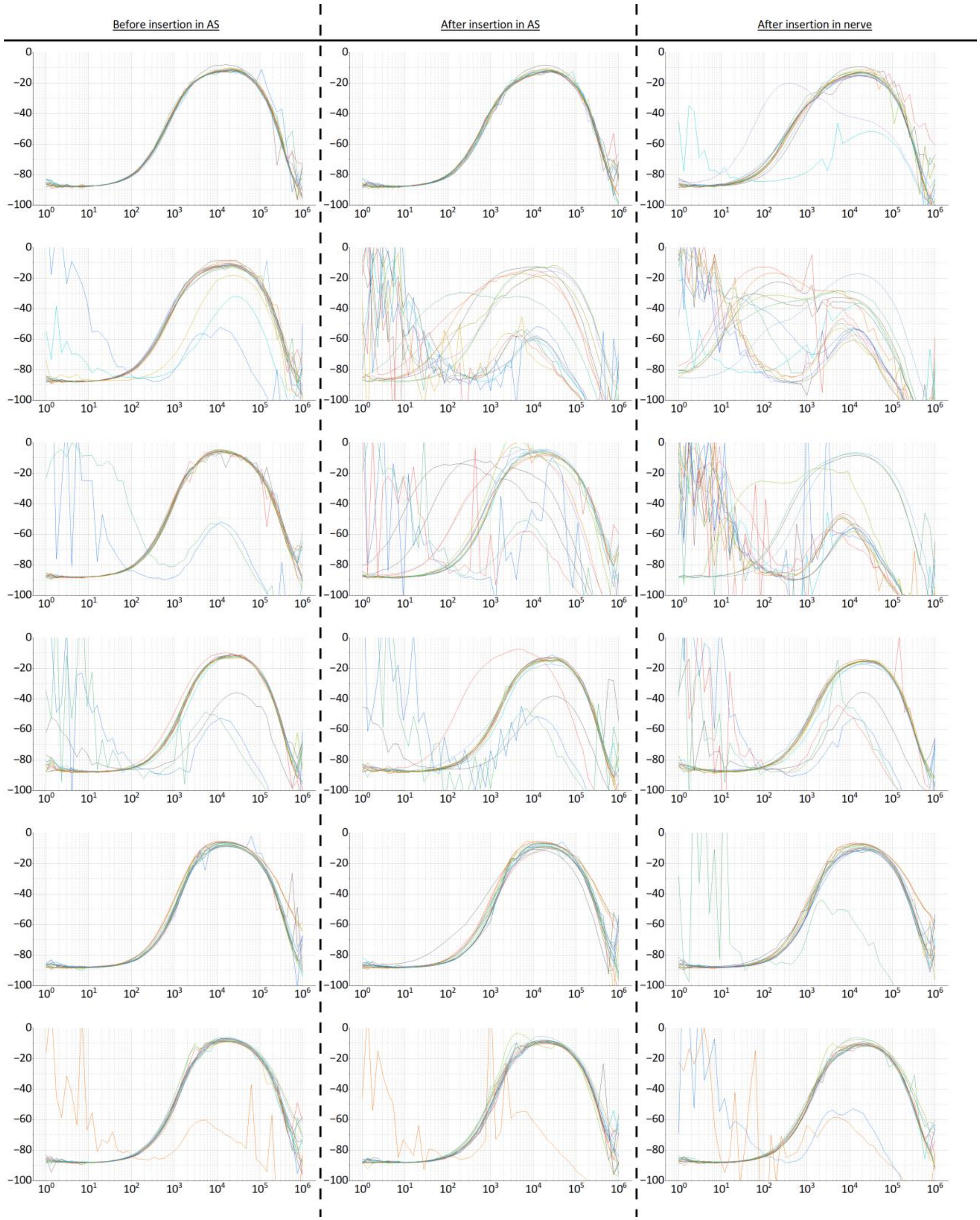


Figure S6: Impedance phases of electrodes from probes used for electrochemical stability tests. X-axes represent frequency [Hz] and y-axes represent phase [°]. Each of the 6 rows belongs to one individual probe. The order of probes is the same as in Figure S5.

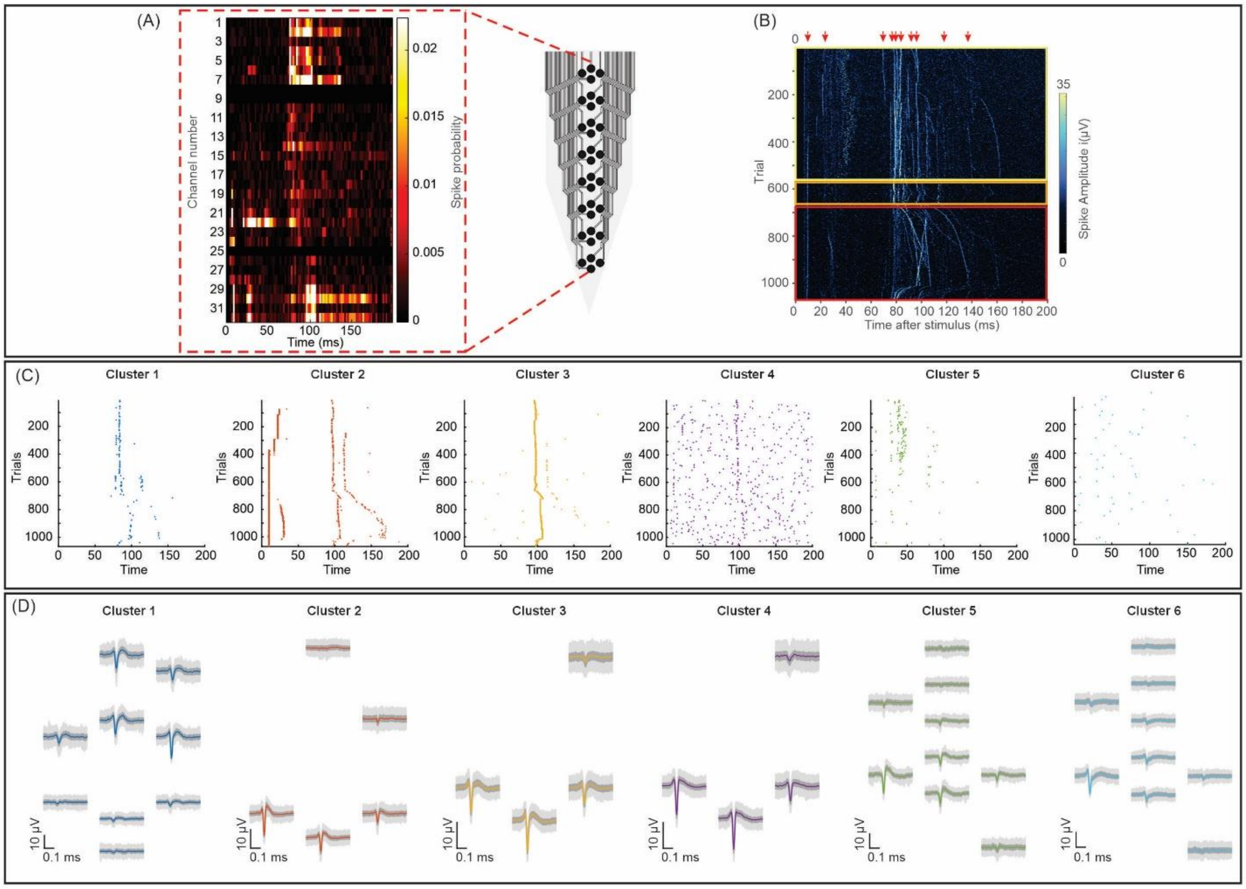

**Figure S7.** ***In vivo*** **characterization of peripheral nerve probes.**
**(A)** Spatial distribution of neural activity across channels highlighted (red dashed rectangular); 32-channel peripheral nerve probe layout. **(B)** A trial-by-trial latency variation heatmap during electrical stimulation shows multiple candidate A-, C-nociceptor/C-non-nociceptor fibers and spontaneous activity on individual recording in vivo. Yellow and orange squares represent low-frequency stimulation protocols, red one high frequency stimulation protocol. **(C)** Functional separation of fibers after manual curation with **(D)** their corresponding waveforms.

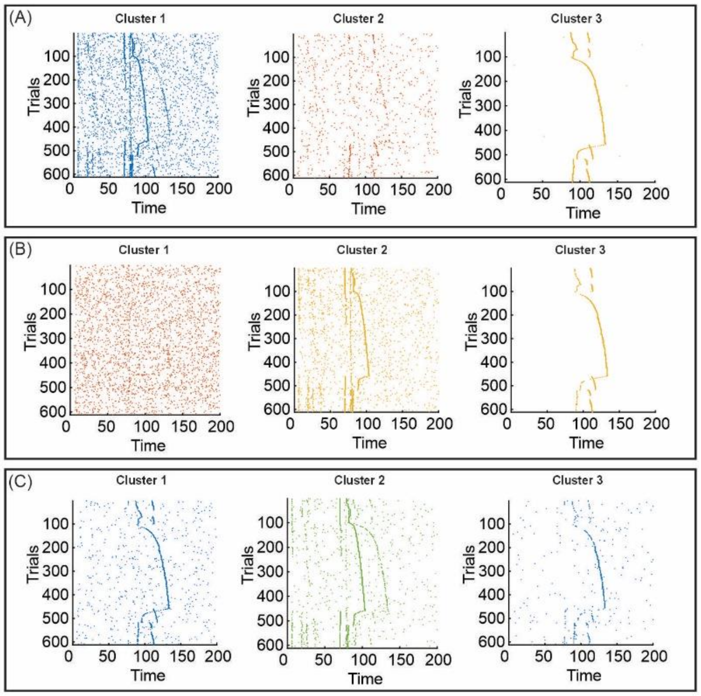


**Figure S8.** Trial-by-trial latency analysis for three best clusters identified by (A) KiloSort4; (B) SpikingCircus2; (C) Tridesclous2.

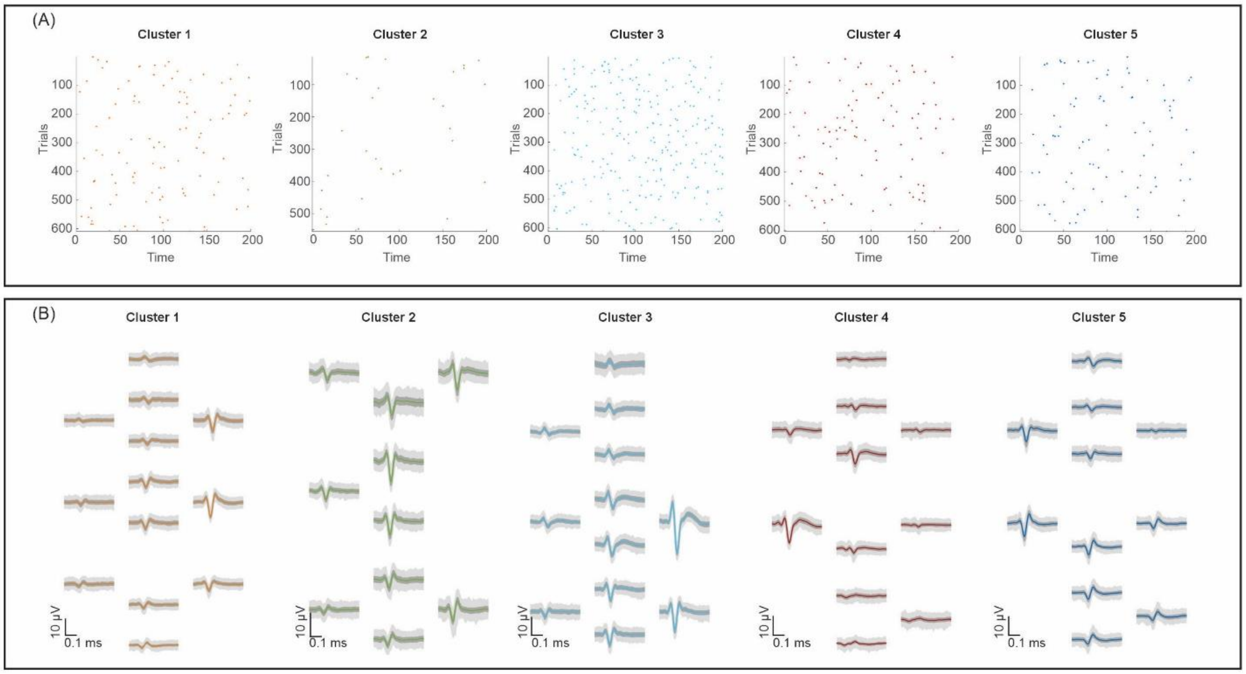


**Figure S9. Non-functional cluster for representative recording. (A)** Functional separation of fibers after manual curation with **(B)** their corresponding waveforms.

**Supplementary Video 1: PNP insertion into saphenous nerve**

In the video, a cannula with a diameter of 900 µm is first inserted through the skin of a euthanzied rat to create a pre-hole. Subsequent to this, a PNP dummy is inserted into the saphenous nerve of the euthanzied rat through the pre-hole. Video shown at 3x real-time speed.
